# A failure of β-amyloid physiological function due to genetic deletion of α7 nicotinic acetylcholine receptors induces an Alzheimer’s disease-like pathology

**DOI:** 10.1101/2021.01.05.425382

**Authors:** Maria Rosaria Tropea, Domenica D. Li Puma, Marcello Melone, Walter Gulisano, Ottavio Arancio, Claudio Grassi, Fiorenzo Conti, Daniela Puzzo

## Abstract

The accumulation of amyloid-beta peptide (Aβ) and the failure of cholinergic transmission are key players in Alzheimer’s disease (AD). However, in the healthy brain, Aβ contributes to synaptic plasticity and memory acting through α7 subtype nicotinic acetylcholine receptors (α7nAChRs). Here, we hypothesized that the α7nAChR deletion blocks Aβ physiological function and promotes a compensatory increase in Aβ levels that, in turn, triggers an AD-like pathology.

To validate this hypothesis, we studied the age-dependent phenotype of α7 knock out mice. We found that α7nAChR deletion caused an impairment of hippocampal synaptic plasticity and memory at 12 months of age, paralleled by an increase of Amyloid Precursor Protein expression and Aβ levels. This was accompanied by other classical AD features such as a hyperphosphorylation of tau at residues Ser 199, Ser 396, Thr 205, a decrease of GSK-3β at Ser 9, the presence of paired helical filaments and neurofibrillary tangles, neuronal loss and astrocytosis.

Our findings suggest that α7nAChR malfunction might precede Aβ and tau pathology, offering a different perspective to interpret the failure of anti-Aβ therapies against AD and to find novel therapeutical approaches aimed at restoring α7nAChRs-mediated Aβ function at the synapse.

## 1. Introduction

Alzheimer’s disease (AD) is the most common neurodegenerative disorder affecting the elderly, but its intricate pathophysiology has prevented the discovery of effective therapies. The Cholinergic and the Amyloid-β (Aβ) Hypotheses represent the two main etiopathological theories proposed to explain the onset and progression of the disease. The Cholinergic Hypothesis (Appel, 1981) has been supported by several evidences indicating that cholinergic transmission is affected in early AD. Indeed, loss of cholinergic neurons in the nucleus basalis of Meynert, decrease of choline acetyltransferase (ChAT) activity and reduction of nicotinic receptors (nAChRs) have been highly correlated with dementia and its progression (Burghaus et al., 2000; Dickson et al., 1995; Engidawork et al., 2001; Kuhn et al., 2015; Mufson et al., 2007; Strada et al., 1992; Whitehouse et al., 1981).

On the other hand, the Aβ Hypothesis (Hardy and Allsop, 1991) posits that the increase and accumulation of Aβ represent the *primum movens* in AD pathophysiology, responsible of synaptic dysfunction triggering downstream events leading to dementia [reviewed in (Gulisano et al., 2018a)].

A third leading actor in this complex picture is tau, a microtubule-associated protein involved in microtubule assembly and stabilization (Wang and Mandelkow, 2015) whose activity and function is regulated by different types of post-translational modifications such as phosphorylation, ubiquitination or glycosylation (Martin et al., 2011). Interestingly, tau shares numerous characteristics with Aβ since both proteins form insoluble deposits, i.e. senile plaques and neurofibrillary tangles (NFTs), respectively (Glenner and Wong, 1984; Grundke-Iqbal et al., 1986), and can aggregate in soluble oligomers whose increase has been highly related to AD severity (Fá et al., 2016; Hölttä et al., 2013; Lasagna-Reeves, 2012; Sengupta et al., 2017). According to the classic Aβ Hypothesis, tau hyperphosphorylation is triggered by Aβ but, recently, it has been demonstrated that the two proteins might act independently or concomitantly to impair synaptic plasticity and memory (Fá et al., 2016; Puzzo et al., 2020), probably converging onto a common molecule represented by amyloid precursor protein (APP) (Puzzo et al., 2017; Wang et al., 2017).

Although pre-clinical and clinical data continue to support the Cholinergic and the Aβ hypotheses, therapeutic strategies aimed at increasing ACh transmission are not able to act as disease-modifying drugs, and approaches expected to cure the disease by decreasing Aβ levels have failed so far. In particular, cholinesterase inhibitors used to treat cognitive symptoms in early to moderate AD patients (Anand and Singh, 2013) do not induce a long-term improvement of cognition and the treatment is not always effective (Connelly et al., 2005; Lemstra et al., 2007). Also, a variety of nAChR agonists, although promising in preclinical studies, had a limited efficacy when experimented in clinical trials, probably for the rapid nAChRs desensitization (Picciotto, 2000). The outcome of clinical trials with anti-Aβ drugs is even more puzzling since the success obtained in animal models of AD has not been replicated in humans. Active and passive immunization against Aβ as well as the use of drugs aimed at preventing Aβ formation have failed, either not showing efficacy or inducing severe side effects [for a review see (Gulisano et al., 2018a)]. Notwithstanding these discouraging results, anti-Aβ therapies are still under investigation with the intent to treat patients in the very early asymptomatic phase of the disease, or to select Aβ-responders based on a personalized approach. However, the latest unsuccessful trials with the anti-Aβ antibody aducanumab and the BACE inhibitor elenbecestat (see www.alzforum.org) are emblematic and confirmed that the Occam’s razor strategy “if Aβ increases in AD patients, clearing Aβ from the brain is the solution” might not be the right choice against AD (Gulisano et al., 2018a; Herrup, 2015; Puzzo et al., 2015).

It is undeniable that AD patients present an increase of Aβ and hyperphosphorylated tau, as well as an impairment of cholinergic transmission (Gulisano et al., 2018a; Ferreira-Vieira et al., 2016; Selkoe and Hardy, 2016). On the other hand, the observation that in the healthy brain Aβ exerts a physiological function mediated by cholinergic receptors (Gulisano et al., 2019) might offer a new perspective to tackle the intricate pathophysiology of AD (Puzzo et al., 2015). Indeed, a variety of studies have demonstrated that Aβ enhances neurotransmitter release (Gulisano et al., 2019; Koppensteiner et al., 2016; Lazarevic et al., 2017) – for a review see (Puzzo et al., 2015) - and facilitates long-term potentiation (LTP) and memory formation (Garcia-Osta and Alberini, 2009; Morley et al., 2010; Palmeri et al., 2017; Puzzo et al., 2011, 2008; Ricciarelli et al., 2014) through α7nAChRs. In fact, picomolar concentrations of Aβ bind α7nAChRs with high affinity (Wang et al., 2000) exerting an agonist-like action that regulates synaptic function (Dineley et al., 2002; Gulisano et al., 2019; Lawrence et al., 2014; Lazarevic et al., 2017; Mura et al., 2012; Oz et al., 2013; Puzzo et al., 2011, 2008). Therefore, a genetic or pharmacological deletion of α7nAChRs prevents the Aβ-induced enhancement of short- and long-term synaptic plasticity as well as memory (Gulisano et al., 2019; Puzzo et al., 2011, 2008).

These observations inspired this work that aimed at understanding whether the Cholinergic and Aβ hypotheses might be unified looking at the disease from a different perspective summarized in one question: what are the consequences of a failure of Aβ physiological function when its endogenous receptor, i.e. α7nAChR, does not work properly?

## 2. Material and Methods

### 2.1 Animals

We used WT (C57BL/6J; RRID:IMSR_JAX:000664) and α7-KO (B6.129S7-Chrna7tm1Bay/J; RRID:IMSR_JAX:003232) purchased from The Jackson Laboratory. Histology was also performed on hippocampal slices from 3×Tg mice (APPSwe, PS1M146V, and tauP301L) genetically engineered by LaFerla and colleagues at the Department of Neurobiology and Behaviour, University of California, Irvine (Oddo et al., 2003). Colonies were established in the animal facilities at University of Catania and Università Cattolica del Sacro Cuore. Housing conditions were controlled maintaining stable hygrometric and thermic conditions (50%; 21°C ± 1°C) on 12 h light/dark cycle with ad libitum access to food and water. All the experiments were performed according to the local Institutional Animal care and Use Committee (approval #327/2013-B, #119-2017-PR, #626-2016-PR) and the European Communities Council Directives (2010/63/EU). Experiments complied with the ARRIVE guidelines and were conducted to minimize animal suffering. To reduce number of animals, we used males for electrophysiological recordings, sex-balanced animals for behavioral experiments, western blotting and ELISA, females for histology and immunohistochemistry. Animals were used at different ages according to our scientific work plan, as detailed in the specific sections.

### 2.2 Electrophysiological field recordings

Extracellular electrophysiological field recordings were performed on 400 mm transverse hippocampal slices as previously described (Gulisano et al., 2019; Puzzo et al., 2017). After cervical dislocation, hippocampi were removed and cut (400 μm thickness) by a manual tissue chopper. Slices were transferred to a recording chamber and perfused (1-2 mL/min) with ACSF (composition in mM: 124.0 NaCl, 4.4 KCl, 1.0 Na_2_HPO_4_, 25.0 NaHCO_3_, 2.0 CaCl_2_, 2.0 MgCl_2_, 10.0 Glucose) kept at 29 °C and continuously bubbled with an O_2_/CO_2_ mixture at 95% and 5%. Slices were allowed to recover for 120 minutes prior to recording. Field excitatory postsynaptic potentials (fEPSPs) were recorded in CA1 *stratum radiatum* by a glass capillary filled with ACSF in response to stimulation of the Schaffer collaterals by a bipolar tungsten electrode. Basal synaptic transmission (BST) was assessed by stimulating with a series of increasing voltage pulses (from 5 to 35 V) to select healthy slices to be used for electrophysiological recordings. For Paired pulse facilitation (PPF) experiments, slices were perfused with the NMDA receptor antagonist (2R)-amino-5-phosphonovaleric acid (APV; 50 μM) for 45 min. Two pulses with a time interval of 10, 20, 30, 40, 50, 100, 200, 500, and 1000 ms were delivered and fEPSP responses were recorded. In another series of experiments, we studied LTP. Baseline was elicited every minute, by stimulating at a voltage able to evoke a response of 35% of the maximum evoked response in BST. After 30-45 min, slices with a stable baseline (slope variation ± 5%) were recorded for 15 min before to induce LTP by a theta-burst (TBS) stimulation, i.e. 3 TBS trains delivered with a 15 seconds inter-train interval with each train consisting in 10 × 100 Hz bursts with 5 pulses per burst with a 200-ms interburst interval, at the test pulse intensity. Recordings were performed and analyzed offline in pClamp 10 (Molecular Devices, Sunnyvale, CA, USA). PPF was plotted as the percentage of the synaptic response of the second against the first delivered stimulus. LTP was plotted as fEPSP (normalized as % of baseline) vs. time (min), or as Residual potentiation, i.e. the average of the last 5 min recording.

### 2.3 Behavioral studies

Fear Conditioning (FC) was performed as previously described (Puzzo et al., 2017). The apparatus consisted in a conditioning chamber, connected to an interface (Kinder Scientific, USA), located in a sound-attenuating box (Campden Inst., UK) with a computer fan installed in one side to provide a background white noise. A webcam mounted on the top of the chamber allowed video recording of the experiment. The floor, made of 36-bar insulated shock grid, was cleaned after each test with 70% ethanol and water. The protocol lasted 3 days. Mice were handled every day for about 5 min before the experiment. During the first day the animal was placed in the conditioning chamber for 2 min prior to the conditioned stimulus (CS) delivery. CS was a tone (2800 Hz and 85 dB) delivered for 30 s. In the last 2 s of the tone, the mouse received a foot shock as an unconditioned stimulus (US) through the electrified grid floor (0.7 mA for 2 s). After the CS/US pairing, the mouse was left into the chamber for 30 s before to be placed in the home cage. Twenty-four hours after training (day 2), the mouse was placed back in the conditioning chamber for 5 min to evaluate contextual fear memory. Forty-eight hours after training (day 3) animals were placed in the conditioning chamber to evaluate cued fear memory. To this end, a novel context was created by using an acrylic black box with a smooth flat floor sprayed with vanilla odorant. After 2 min (pre-CS test), the mouse was exposed to the same tone used during the training for 3 min (CS test). Freezing (absence of movement except for that needed for breathing) was manually scored during the three days by two different operators and the averaged value was used to perform the analyses.

Novel Object Recognition (NOR) was performed as previously described (Gulisano et al., 2018b). The arena was a plastic white box (50 x 35 x 45 cm) placed on a lab bench. A webcam, connected to the computer, was fixed on the wall. The NOR protocol was performed in 5 days: 3 days of habituation, 1 day of training (T1) and 1 day of testing (T2). Objects were designed by a computer aided design software (Solidworks, France) and printed in polylactic acid with a Prusa i3-inspired 3D printer of our design. After each trial, the box and the objects were cleaned with 70% ethanol and dried with absorbent paper. During the first day (habituation to the arena), the mouse was put into the empty arena and allowed to explore it for 10 min. During the second and the third day (familiarization with objects), the mouse was put into the arena containing two different objects, randomly chosen among our object collection and changed from day to day, for 10 min. During the fourth day, NOR training session (T1) was performed. The mouse was put into the arena and allowed to explore for 10 min two identical objects placed in the central part of the box, equally distant from the perimeter and the center. During the fifth day (24 h after T1), the mouse underwent the second trial (T2) to test memory retention for 10 min. Mice were presented with two different objects, respectively a “familiar” (i.e. the one used for T1) and a “novel” object. For Novel Object Location (NOL), the mouse was put into the arena and allowed to explore for 10 min two identical objects placed in one side of the box. On the day after, the location of one object was changed and the mouse underwent the T2 for 10 min. For both NOR and NOL experiments, animal exploration - defined as the mouse pointing its nose toward the object from a distance not > 2 cm - was measured in T2. We analyzed: i) percentage exploration of familiar vs. novel object; ii) discrimination (D) index, “exploration of novel object minus exploration of familiar object/total exploration time”; and iii) total exploration time. Mice with a total exploration time < 5 s were excluded from analysis.

### 2.4 Determination of Aβ levels

Briefly, hippocampal tissues from 12M α7 KO and WT mice were sonicated in lysis buffer (10 μl/mg tissue) containing 5M guanidine- HCl/50mM Tris, pH 8.0). Sonicates were then diluted ten- fold with Dulbecco’s PBS containing 1× protease inhibitor cocktail (Sigma). Levels of murine Aβ (1–42) were measured by enzyme-linked immunosorbent assay (ELISA) using commercial kits (Thermo Fisher Scientific, cat# KMB3441) following manufacturer’s instructions. All assays were performed on F-bottom 96-well plates (Nunc, Wiesbaden, Germany). Tertiary antibodies were conjugated to horseradish peroxidase. Wells were developed with tetramethylbenzidine and measured at 450 nm.

### 2.5 Western Blotting on hippocampal homogenates

Western blot (WB) analysis was performed as previously described (Gulisano et al., 2019; Li Puma et al., 2019) with minor modifications. Whole hippocampi from 9 and 12 months-old WT and α7 KO mice were homogenized in RIPA buffer (Thermoscientific) in the presence of phosphatase and protease inhibitors (Thermoscientific), and sonicated 3 times for 10 minutes on ice. Protein concentrations were determined by Bradford protein assay (Biorad) and 40μg of total proteins were then loaded onto 4-15% Tris-glycine polyacrylamide gels (Biorad) for electrophoretic separation and then transferred onto 0.45 or 0.22 μm nitrocellulose membranes (Amersham Biosciences, Buckinghamshire,UK). Membranes were blocked for 1-hour, at RT, in either a solution of 5% nonfat dry milk in Tris-buffered saline containing 0.1% Tween-20 before incubation overnight at 4°C with the following primary antibodies: mouse 4G8, that recognizes residues 17–24 of Aβ and the same sequence in APP full length (BioLegend San Diego, California, USA; 1:1000); mouse C-term APP Y188 (Abcam; Cambridge UK; 1:1000); mouse M3.2 (BioLegend; 1:1000), that recognizes residues 10-15 of murine Aβ and the same sequence in APP full length; mouse pTau Ser199 (Thermo fisher Scientific; 1:1000); rabbit pTau Ser396 (SAB; Signalway Antibody Co., Ltd.; 1:1000); rabbit pTau Thr205 (SAB; 1:1000); rabbit pGSK-3β Ser9 (Cell Signaling; 1:1000). Mouse anti-GAPDH (Abcam; 1:5000), rabbit total GSK-3β (Cell signaling; 1:1000) and mouse Tau-5 (Thermo fisher Scientific; 1:1000) were used as loading controls. After incubation with HRP-conjugated secondary antibodies (Cell Signaling Technology) visualization was performed with ECL plus (Amersham Biosciences) using UVItec Cambridge Alliance. Molecular weights for immunoblot analysis were determined using Precision Plus Dual Color Standards (Biorad).

### 2.6 Tissue preparation for microscopy studies

For histology, NeuN and GFAP immunohistochemistry, fresh brains were removed, immersed in 10% formalin for 72 hrs and then transferred in 4% PFA in phosphate buffer until use. For PHF-1 immunohistochemistry, animals were anesthetized by intraperitoneal injection of a cocktail of Zolazepam plus Tiletamine (120 mg/Kg) and Medetomidine (80 μg/Kg), and perfused through the ascending aorta with a flush of physiological saline followed by 4% paraformaldehyde. Brains were removed and post-fixed in 4% PFA in phosphate buffer for 5 days.

Sections (50 μm thickness) were sequentially cut with a Vibratome within 2.350 and 1.725 mm lateral range and processed for immunohistochemistry or histological stains.

### 2.7 Congo Red stain

Congo Red stain was performed as previously described (Wilcock et al., 2006). Sections were incubated for 20 min in a fresh prepared alkaline saturated NaCl solution. Briefly, NaCl was added to an 80% ethanol solution while stirring, until the formation of an undissolved NaCl layer (about 5 mm thick) and 1% NaOH 1 M was added before use. Sections were then incubated in 0.2 % Congo Red solution (Sigma-Aldrich) for 30 min and rinsed five times in 95% ethanol.

### 2.8 Bielschowsky stain

Bielschowsky silver stain was performed as suggested by the manufacturer (Bielschowsky silver stain kit, VitroVivo Biotech, cat.#VB-3015). Briefly, sections were incubated in pre-warmed 40°C Silver Nitrate solution for 18 minutes, washed in distilled water and then incubated with silver ammonium solution at 40°C for 30 minutes. Subsequently, slices were placed in the developing solution for 75 seconds and in 1% ammonium hydroxide solution for 60 seconds. Slices were then washed in distilled water and incubated in 5% sodium thiosulfate solution for 5 minutes.

### 2.9 Immunohistochemistry

#### Antibodies

The following primary antibodies were used: mouse anti-PHF-1 (1:50) detecting tau Ser396/Ser404 phosphorylation sites (generous gift of Dr. Peter Davies); mouse anti-NeuN (1:100; Millipore, cat.#MAB377); mouse anti-GFAP (1:350; Sigma-Aldrich, cat.#G3893).

#### Immunofluorescence

Immunofluorescence was performed as previously described (Melone et al., 2019). Sections were incubated in 10% NDS (1 hr), followed by a solution containing PHF-1 primary antibody overnight at RT. Sections were rinsed in Tris (0.5 M buffered saline, pH 7.4) incubated first in 10% NDS (15 min), and then in a solution containing the secondary antibody (Alexa Fluor 488, 1:230; Jackson) for 2 hrs at RT. Sections were washed in Tris, mounted, and coverslipped with propyl-gallate 0.1M in glycerol-PBS solution (9:1). Omission of the primary antibodies in sections from each experimental group resulted in a lack of specific staining in the corresponding channel (negative control). To quench lipofuscin autofluorescence, at the end of immunofluorescence protocols, sections were incubated for 5 min in 0.1% Sudan Black dissolved in 70% alcohol. To minimize procedural variability, sections from all experimental groups, were exposed to immunofluorescence procedure in parallel.

#### Immunoperoxidase

Immunoperoxidase was performed as previously described (Melone et al., 2019). For both NeuN and GFAP optimal detection (i.e. immunoreactivity against the background level) was determined by testing different conditions (e.g., inclusion or exclusion of detergents, and a range of dilutions for primary antibodies) in a set of pilot trials. Sections were treated with H_2_O_2_ (1% in Tris for 25 min) to remove endogenous peroxidase activity, rinsed in Tris and then incubated in 10% NGS (1 hr) followed by a solution containing NeuN or GFAP primary antibodies (overnight at RT). Sections were rinsed in Tris and incubated first in 10% NGS (15 min) and then in a solution containing biotinylated secondary antibody (1:180; Jackson) for 2 hrs at RT. Subsequently, they were rinsed in Tris, incubated in avidin-biotin peroxidase complex (ABC Elite PK6100, Vector), washed in Tris, and incubated in 3,3’diaminobenzidine tetrahydrochloride (DAB; 0.05% in 0.05 M Tris buffer, pH 7.6 with 0.03% H_2_O_2_). Then, sections were washed, mounted, and coverslipped with dpx mounting medium. Method specificity was assessed by substituting NeuN and GFAP primary antibodies with Tris, resulting in the absence of immunoreactivity. To minimize procedural variability, sections from all experimental groups, were exposed to immunoperoxidase procedure in parallel.

#### Data collection and analysis

For confocal microscopy, PHF-1 immunolabeled sections were scanned with a Leica SP2 TCS-SL microscope. For 20x of CA1, microscopical fields were acquired as 512 x 512 pixel images (pixel size of 750 nm) with pinhole 1.4 Airy unit, and to improve signal/noise ratio, 4 frames of each image were averaged. For quantitative microscopy of stratum oriens (so), stratum pyramidalis (sp), and stratum radiatum (sr), microscopical fields were acquired as 512 x 512 pixel images (pixel size of 465 nm) with a planapo 63 × objective (numerical aperture 1.4) and pinhole 1.0 Airy unit. To improve signal/noise ratio, 4 frames of each image were averaged. CA1 microscopical fields were randomly selected (8 microscopical fields/layer/4 sections from 2 animals for each experimental group). To avoid the influence of the acquisition parameters (i.e. photomultiplier gain and offset) on fluorescence intensity, all microscopical fields from all conditions were scanned and acquired with the same setting. As previously described (de Vivo, 2010), photomultiplier gain and offset were set so that the brightest pixel was just slightly below saturation, and the offset such that the darkest pixels were just above zero. To avoid the effects of the surface-depth gradient on immunodetection (Melone et al., 2005), all microscopical fields were acquired at a z-axis level yielding the maximum brightness of immunopositive profiles (de Vivo, 2010; Melone et al., 2005).

For quantitation of intensity, randomly selected subfields of 32 × 32 μm from the original microscopic fields (Melone et al., 2019) (32 for each experimental group/layer; with a total of 96 subfields for each condition) were used.

Optimal visualization of immunoreactivity was achieved by setting the threshold value to the mean pixel value over the field under study in WT and α7 KO groups (Melone et al., 2005). Intensity of threshold subfields was calculated by Image J (Schneider et al., 2012). Number and mean size of immunoreactive puncta of each subfield were obtained by transforming images to binary mode and calculated using Image J (Bozdagi et al., 2000; Bragina et al., 2006; Schneider et al., 2012). For PFH-1 positive neuronal-like cells, intensity of 2-4 regions of interested (ROI) within the cytoplasm, was calculated (area between 30 and 40 μm²) (de Vivo, 2010). For the intensity of PHF-1 positive dendritic-like profiles, value was extracted by plotting intensity pixel values along the major axis of profiles using Image J (Melone et al., 2019).

For light microscopy studies, hippocampus and CA1 of Congo Red and Bielschowsky stained sections were acquired at 4 × and 20 × and at 4 × and 40 × original magnifications, respectively (4-6 sections/2 animals for each experimental group). For NeuN and GFAP studies, hippocampi of immunostained sections were acquired at 4 ×, and at 40 × for quantitative studies in so, sp, sr. NeuN and GFAP positive cells were manually identified, and to estimate the density of cells, the area of microscopical fields (18 microscopical fields/layer/6 sections from 3 animals for each experimental group; with a total of 54 fields for each condition) was calculated by Image J.

### 2.10 Statistics

All experiments were performed by researchers blind with respect to treatment. All data were expressed as mean ± standard error mean (SEM). Statistical analysis was performed by using different tests, based on preliminary analyses of normal distribution. ANOVA for repeated measures was used to analyze PPF and LTP (120 minutes of recording after tetanus). One-way ANOVA with Bonferroni’s post-hoc correction was used for PPF single time intervals and LTP graphs displaying residual potentiation (average of the last five minutes of LTP recording). Two-tailed t-test was used for analyses of behavioral parameters, NeuN and GFAP immunohistochemistry. One sample t-test was used to compare D with zero in NOR and NOL. Given the non-normal distribution of data, assessed by D’Agostino & Pearson normality, we used Mann-Whitney test for ELISA and PHF-1 immunoreactivity; Mann-Whitney and Kruskal-Wallis One Way tests for WB experiments. Systat 9, Graphpad Prism 8, and Sigmaplot 14 software were used for statistical analyses. The level of significance was set at P < 0.05.

## 3. Results

### 3.1 Synaptic plasticity and memory are impaired in 12-month-old α7 KO mice

The role of α7nAChRs in cognitive functions (Picciotto, 2000) relies on their ability to modulate synaptic function through the regulation of glutamate release (Cheng and Yakel, 2015). Thus, we first evaluated whether their genetic deletion affected paired-pulse facilitation (PPF), a form of short-term plasticity that might reflect release probability. PPF resulted unchanged in slices from α7 KO mice at 3, 6 and 9 months of age, but was increased at 12 months (M) compared to slices from age-matched WT animals (Figure 1A). We then investigated long-term plasticity by recording LTP at hippocampal CA3-CA1 synapses. Potentiation was normal in hippocampal slices from animals at 3 and 6M but decreased with age (Figure 1B), especially at 12M, in slices from α7 KO mice (Figure 1B, 1C).

**Figure 1.**
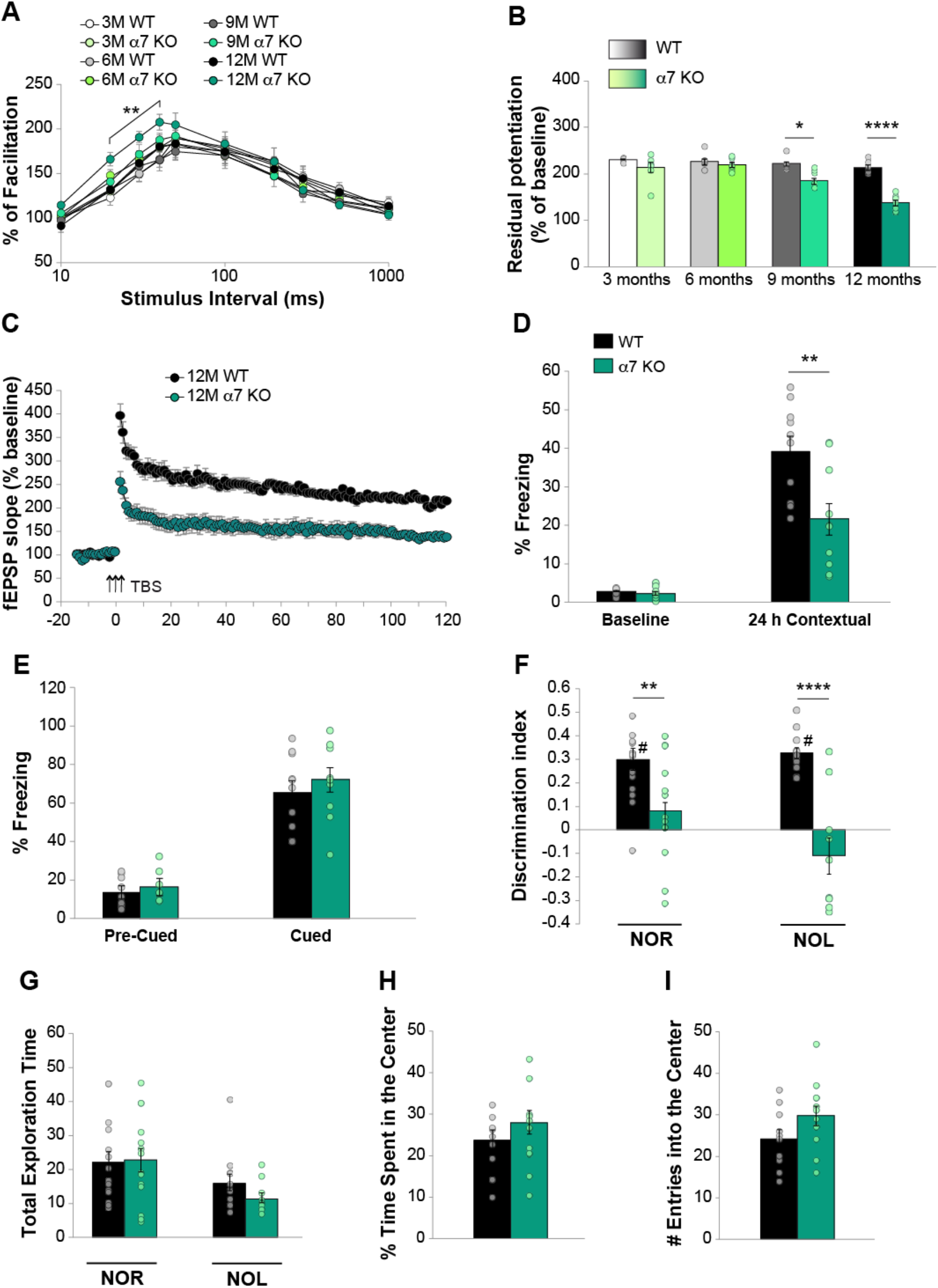
Synaptic plasticity and memory are impaired in 12-month-old α7 KO mice. **A)** Age-dependent modification of PPF (P = 0.010 in 12M α7 KO vs. WT; N = 9 slices from 6-7 3-month-old animals; N = 7 slices from 5-6 animals at other ages). **B)** Age-dependent modification of LTP residual potentiation (averaged last 5 recording points) (Bonferroni’s P = 0.027 and P < 0.0001 between α7 KO and WT at 9 and 12 M. N = 7 slices from 5-6 animals for each condition). **C)** Comparison of LTP curves in slices from 12 M α7 KO and WT controls (F_(1,12)_ = 35.1837, P < 0.0001). TBS = theta-burst stimulation. **D)** Contextual fear memory was impaired in α7 KO mice (t_(18)_ = 3.06, P = 0.007; N = 10/10. **E)** No differences were detected in cued fear memory (t_(18)_ = 0.722, P = 0.479). **F)** Discrimination index (D) was impaired in α7 KO mice (NOR: t_(27)_ = 2.92, P = 0.007; N = 15 WT/12 α7 KO; NOL: t_(27)_ = 5.882, P < 0.0001; N = 12 WT/10 α7 KO). WT but not α7 KO mice were able to learn (P < 0.0001 D vs. zero). **G)** Total exploration time was similar among genotypes (NOR: t_(27)_ = 0.143; P = 0.887; NOL: t_(20)_ = 0.543; P = 0.887). **H-I)** No differences were present in exploratory and anxiety-like behavior tested by the open field task (t_(20)_ = 1.081, P = 0.293 for % time spent into the center; t_(20)_ = 1.632, P = 0.118 for number of entries into the center; N = 10 WT/12 α7 KO). *P < 0.05, **P < 0.01, ****P < 0.0001, #P ≠ 0. Data expressed as mean ± SEM.

The observation that release probability and plasticity were impaired in 12M α7 KO mice in an age-dependent manner, prompted us to study different types of memory known to be impaired in AD (Puzzo et al., 2014a). Evaluation of contextual fear memory 24 hrs after training showed that the amount of freezing behavior was impaired in α7 KO compared to WT controls (Figure 1D), whereas no differences were found in amygdala-dependent cued fear memory (Figure 1E).

We then evaluated recognition and spatial memory through Novel Object Recognition (NOR) and Novel Object Location (NOL) tasks. Analyses of Discrimination index (D = exploration of novel object minus exploration of familiar object/total exploration time) showed an impairment of memory in α7 KO (Figure 1F). Comparison of D with zero confirmed that only WT mice were able to discriminate between the old and the novel object or its different spatial location. No differences were detected in total exploration time between the two groups of mice (Figure 1G).

Open field test showed that locomotor activity and anxiety-like behavior were not affected (Figure 1H,I), suggesting that the impairment of memory found in α7 KO was not due to motor or motivational defects.

Thus, α7 KO mice showed an age-dependent impairment of hippocampal synaptic plasticity and memory.

### 3.2 The lack of endogenous α7nAChRs induces an increase of Aβ production and APP expression

The impairment of synaptic plasticity and memory found in α7 KO mice might be due to the sole alteration of cholinergic transmission. However, this scenario is also compatible with the hypothesis that the cognitive phenotype is triggered by the failure of Aβ-mediated synaptic homeostasis. Indeed, in a situation in which Aβ is not able to adequately exert its physiological functions through α7nAChRs, a feedback mechanism might occur, inducing a compensatory increase of Aβ production. High levels of Aβ might in turn be responsible for the impairment of synaptic plasticity and memory found in α7 KO mice. To test this hypothesis, we evaluated whether the age-dependent damage of LTP and memory was paralleled by changes in Aβ production. We performed the enzyme-linked immunosorbent assay (ELISA) for mouse Aβ_42_ on hippocampal homogenates from 12M α7 KO and WT mice and confirmed that the deletion of α7nAChR induced a significant increase of the peptide levels (Figure 2A). Because Aβ is produced by APP cleavage, we next verified whether this feedback mechanism acted through a modification of APP expression. We found an increase of APP full-length expression in 12M α7 KO hippocampi (Figure 2B,C) that was confirmed by 3 different antibodies: Y188, 4G8, and M3.2. Furthermore, 4G8 (recognizing both human and murine Aβ) and M3.2 (specific for murine Aβ) allowed detecting a 24 KDa band, presumably corresponding to soluble aggregates (i.e., pentamers), that was significantly increased in α7 KO hippocampi (Figure 2D,E). However, Congo Red staining did not reveal the presence of hippocampal senile plaques (Figure 2F).

**Figure 2.**
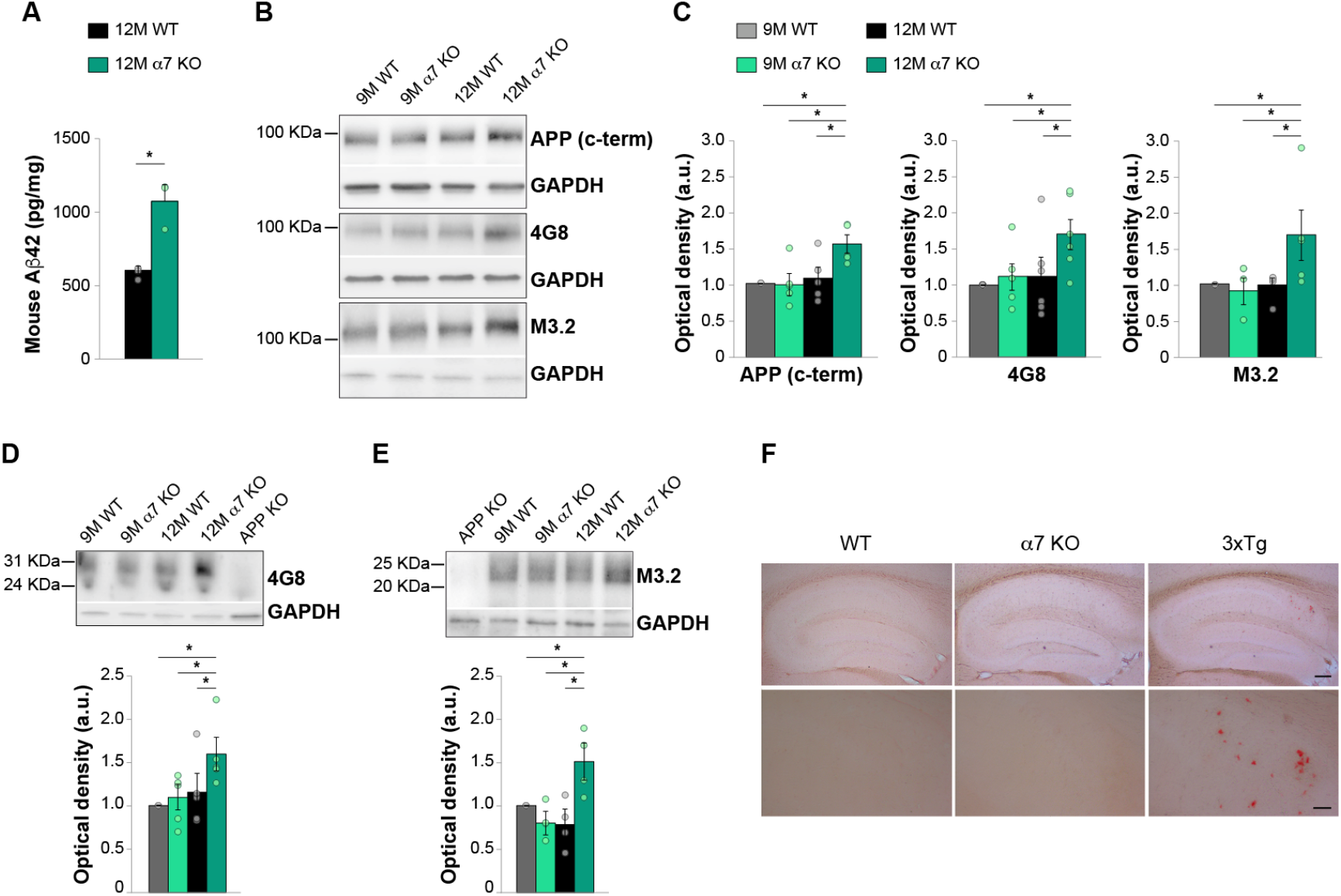
Aβ levels and APP expression increase in 12M α7 KO mice. **A)** ELISA revealed an increase of Aβ42 levels in hippocampal homogenates from α7 KO compared with WT mice (Mann-Whitney Rank Sum Test, P = 0.036, N = 3 and 4 animals, respectively). **B)** Western blot analysis comparing the expression of APP in hippocampi from WT and α7 KO animals at 9M and 12M assessed by 3 different antibodies, i.e. 4G8, Y188 and M3.2. **C)** Bar graphs showing the results of the densitometric analysis of the Western blots reported in (B) indicated a significant increase of APP expression in hippocampi from 12M α7 KO mice compared to WT (Kruskal-Wallis One Way Analysis of Variance on Ranks, P = 0.038 for Y188, N = 5 for each condition; P = 0.040 for 4G8, N = 6 for each condition and P = 0.046 for M3.2; N = 5 for each condition). GAPDH expression level was used as loading control here, in D and E. **D)** An increase of a ≈ 25 kDa band, presumably representing Aβ oligomers, was detected in 12M α7 KO hippocampi either when using the 4G8 (Kruskal-Wallis One Way Analysis of Variance on Ranks, P = 0.035, N = 5 for each condition) or **E)** the M3.2 antibody, specific for murine Aβ (Kruskal-Wallis One Way Analysis of Variance on Ranks; P = 0.030, N = 4 for each consition). APP KO mice were used as negative controls either in D and E. **F)** No plaques were detected using Congo red staining in α7 KO hippocampal slices. Hippocampal slices from 3×Tg mice are used as a positive control. Upper panels: 4 × magnification, scale bar 100μm; Lower panels: 20 × magnification, scale bar 50 μm. *P < 0.05. Data expressed as mean ± SEM.

Overall, these findings suggest that the absence of α7nAChRs triggers an age-dependent increase of soluble Aβ and APP, which parallels the impairment of synaptic plasticity and memory, with no plaque formation.

### 3.3 The lack of endogenous α7nAChRs triggers tau hyperphosphorylation through GSK-3β modulation

Tau hyperphosphorylation and accumulation of paired helical filaments (PHF) leading to NFTs formation represent hallmarks of AD brain lesion highly correlated with cognitive impairment (Iqbal et al., 2005). Here, we first investigated whether α7nAChRs deletion modified tau phosphorylation at different residues known to be associated with neurodegeneration and AD, i.e., Ser 199, Ser 396, and Thr 205 (De Chiara et al., 2019; Neddens et al., 2018). Western blotting analysis showed a significant increase of pTau expression at Ser 199 and Ser 396 (Figure 3A), and Thr 205 (Figure 3B) in hippocampi from 12M α7 KO mice compared to WT.

**Figure 3.**
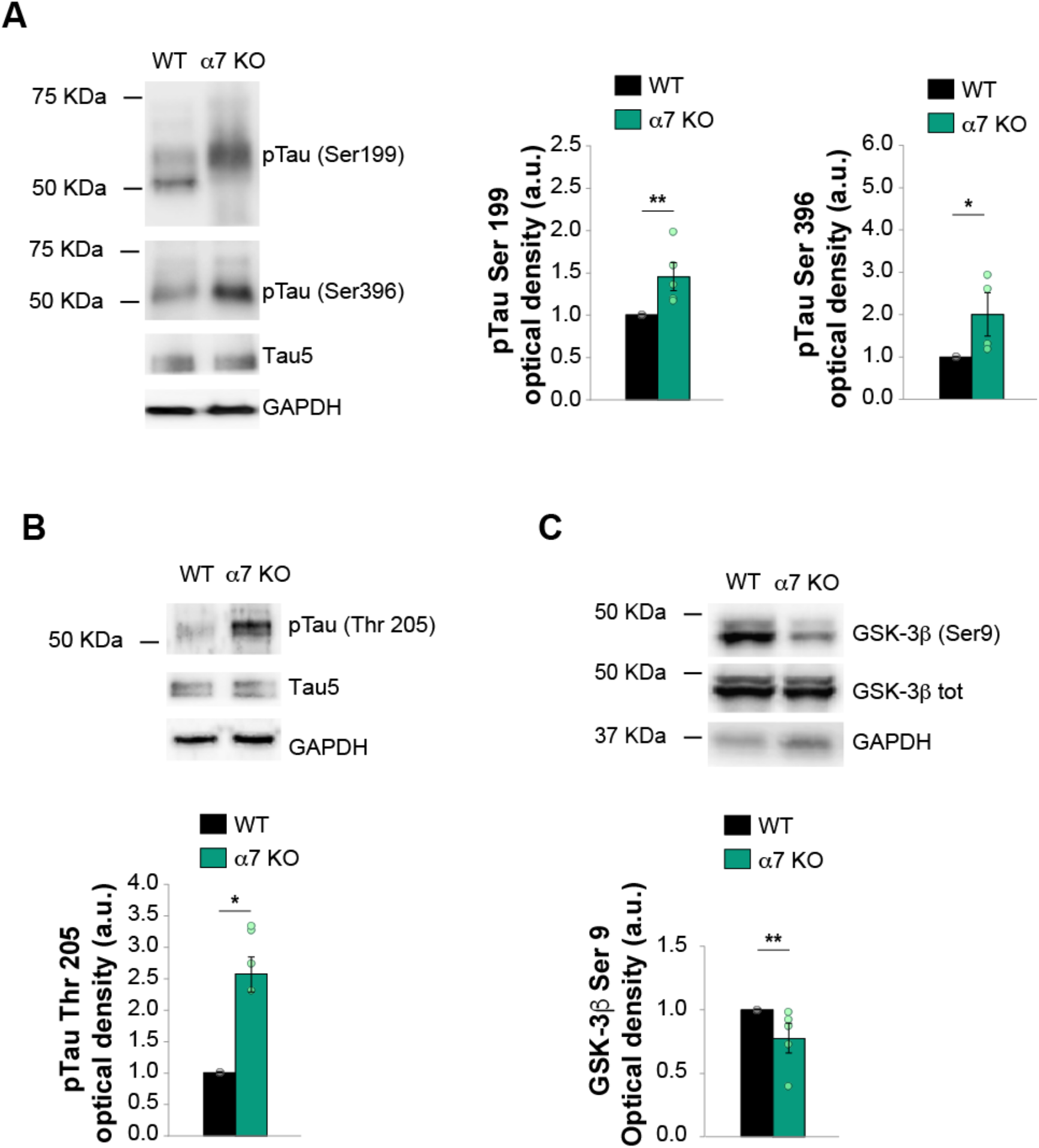
Tau is hyperphosphorylated and GSK-3β dysregulated in hippocampi from 12M α7 KO mice. **A)** Representative images of WB assay (cropped images based on MW here and in the following panels) for the expression of phosphorylated tau (pTau) at Ser 199 and at Ser 396 performed on hippocampi from 12M WT and α7 KO mice. On the right, bar graphs showing the increase of pTau Ser 199 expression in hippocampi from α7 KO compared to WB (Mann-Whitney Rank Sum Test for pTau Ser 199: P = 0.008; N = 5/5; for pTau Ser 396: P = 0.028; N = 4/4). Tau5, specific for murine tau, was used to normalize pTau densitometric signal from WT and α7 KO tissues. GAPDH was used as loading control. **B)** WB essay for the expression of pTau at Thr 205. Lower panel, bar graph showing the increase of the expression of pTau at Thr 205 in hippocampi of α7 KO compared to WT (Mann-Whitney Rank Sum Test, P= 0.028; N = 4/4). **C)** WB images showing the expression of GSK-3β phosphorylated at Ser9, total GSK-3β and GAPDH as loading control. Lower panel, bar graph showing the decrease of GSK-3β Ser 9 in α7 KO hippocampi compared to WT (Mann-Whitney Rank Sum Test; P = 0.007; N = 5/5). *P < 0.05, **P < 0.01. Data expressed as mean ± SEM.

We then turned our attention onto glycogen synthase kinase-3β (GSK-3β), considered a crucial molecule in AD, being a possible molecular link between Aβ and tau pathology (Llorens-MarÃ-tin et al., 2014). In particular, we focused on GSK-3β auto-inhibitory phosphorylation site on Ser 9 whose dysregulation induces an abnormal activation of GSK-3β leading, in turn, to tau hyperphosphorylation (Hanger and Noble, 2011). We found a decrease in Ser 9 phosphorylation of GSK-3β in hippocampi from α7 KO mice (Figure 3C) that paralleled the increase of tau phosphorylation.

### 3.4 The lack of endogenous α7nAChRs causes tau accumulation and deposition in neurofibrillary tangles

We then investigated the presence of PHFs in hippocampal slices, suggestive for a more advanced stage of pathology. We used a PHF-1 antibody (a generous gift of Dr. Peter Davies) that detects Ser396/Ser404 phosphorylation sites, known to be associated with NFT formation (Götz et al., 2001).

Confocal microscopy revealed an increased PHF-1 immunoreactivity (IR) in the CA1 area of 12M α7 KO hippocampi compared to WT (Figure 4A), particularly evident in the CA1 *stratum pyramidalis* and *stratum radiatum* (Figure 4B). Analyses of PHF-1 IR revealed an increase of mean size positive puncta in the *stratum oriens*, neuronal-like cells in the *stratum pyramidalis*, and dendritic-like profiles in the *stratum radiatum* (Figure 4C-E). Consistently, Bielschowsky silver staining detected an increase of intraneuronal NFTs in the hippocampus and neocortex from α7 KO (Figure 4F).

**Figure 4.**
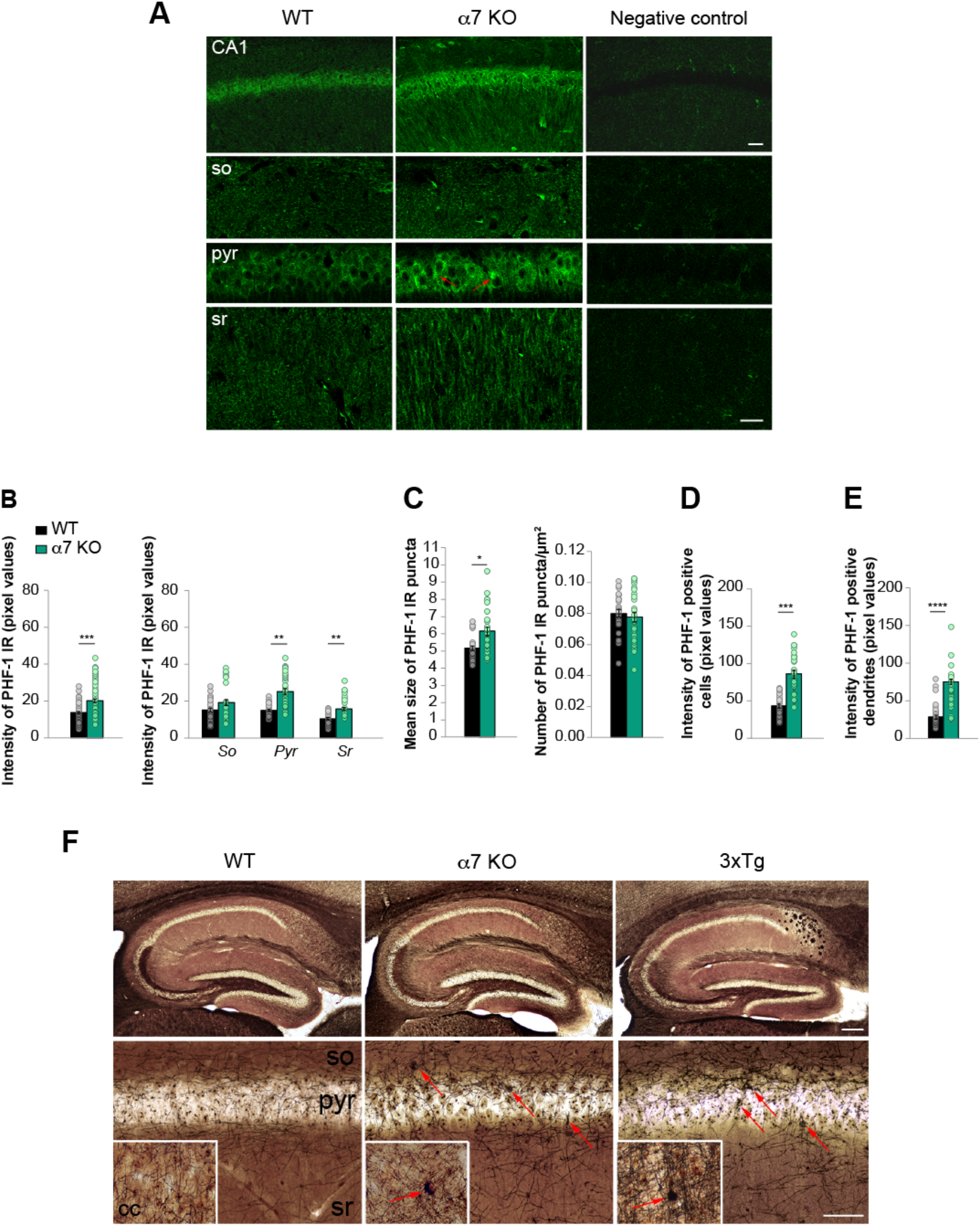
PHFs immunoreactivity and neurofibrillary tangles in hippocampi from α7 KO mice. **A)** Representative confocal images of PHF-1 immunofluorescence in the hippocampus from 12-15M WT and α7 KO mice. Negative control represents WT sections treated with the same immunofluorescence protocol, omitting the primary antibody. Upper panels: CA1 area, 20 ×, scale bar 50 μm; Lower panels: stratum oriens (so), stratum pyramidalis (pyr) and stratum radiatum (sr), 60 ×, scale bar 25 μm. Red arrows indicate intraneuronal accumulations. **B)** Bar graphs showing the increase of PHF-1 immunoreactivity (IR) in hippocampi from α7 KO (Mann-Whitney test P = 0.0002). On the right, analyses of PHF-1 IR in so, pyr and sr. **C)** Bar graphs showing puncta staining analyses in so. Positive puncta mean size increased in α7 KO (P = 0.0113), whereas number of puncta was not modified (P > 0.99). **D)** Bar graphs showing analyses of cytoplasmic PHF-1 IR intensity in the pyr. Positive cells increased in α7 KO (P = 0.0002). **E)** Bar graphs showing an increase of PHF-1 IR intensity in apical dendrites of the sr in α7 KO (P < 0.0001). N = 32 subfields (4 sections from 2 animals/genotype). **F)** Bielschowsky silver staining showed the presence of silver-positive NFTs in α7 KO. Hippocampi from 3×Tg mice were used as positive control. Representative inserts showing a clear identifiable NFT in α7 KO and 3×Tg cerebral cortex (cc). Red arrows indicate intraneuronal accumulation. Upper panels: 4 ×, scale bar 100 μm; Lower panels: CA1 area, 40 ×, scale bar 50 μm; Inserts: 40 ×, scale bar 50 μm. *P < 0.05, **P < 0.01, ***P < 0.001, ****P < 0.0001. Data expressed as mean ± SEM.

Overall, these data indicated that α7 KO mice present a dysregulation of tau phosphorylation resulting in an increase of PHFs and NFTs.

### 3.5 The lack of endogenous α7nAChRs induces neuronal loss and astrocytosis

Besides Aβ and tau pathology, because hippocampal neuronal loss highly correlates with cognitive deficits and AD progression (Donev et al., 2009), we evaluated whether α7 KO mice presented neuronal depletion. We found a decrease in the mean number of NeuN positive cells in hippocampal slices from α7 KO compared to WT controls (Figure 5A). A statistically significant decrease was found in the *stratum pyramidalis* and *stratum radiatum*, whereas no differences were detected in the *stratum oriens* (Figure 5A).

**Figure 5.**
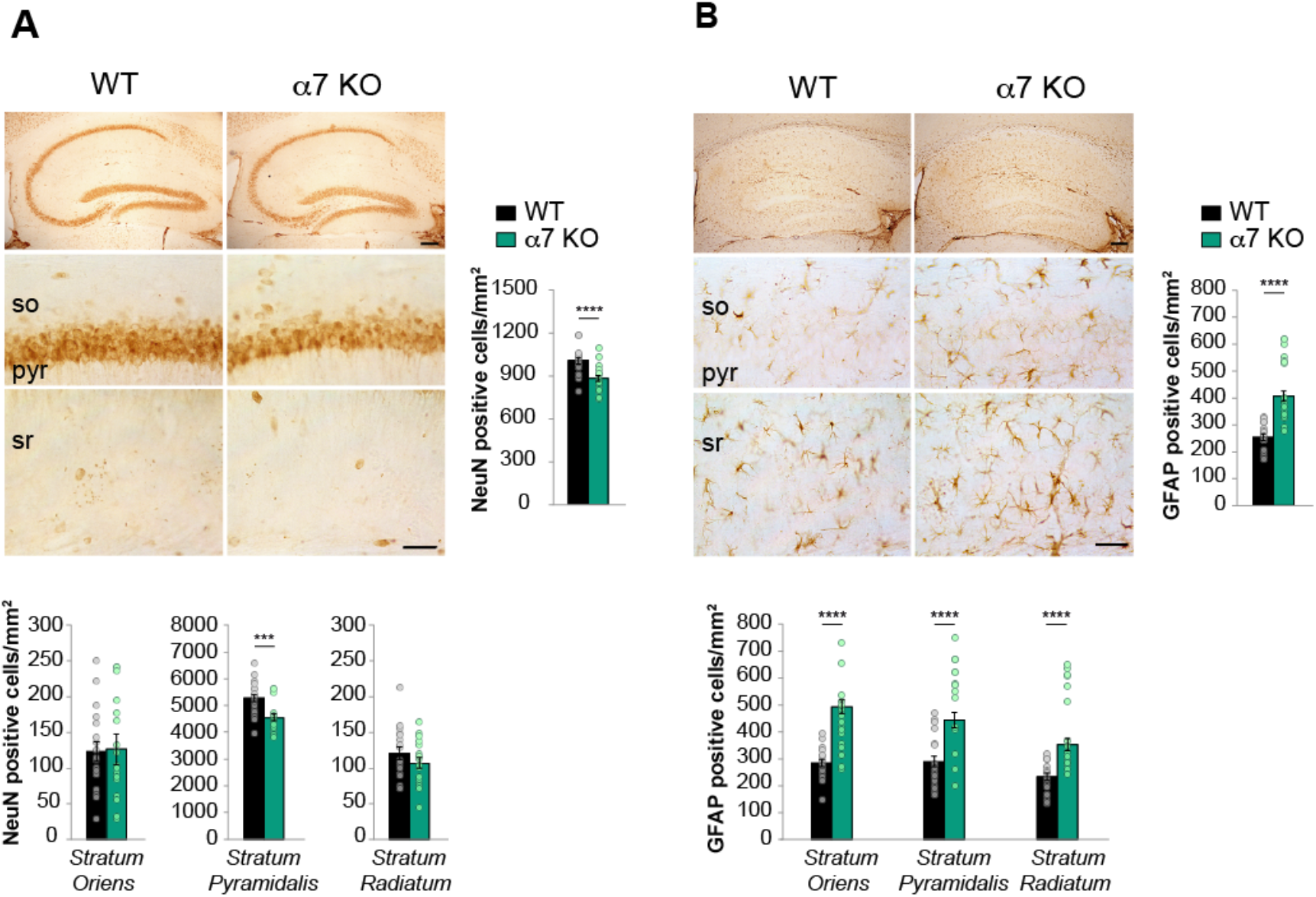
Neuronal loss and astrocytosis increase in α7 KO hippocampi. **A)** Representative images of NeuN staining in the hippocampal formation of 12-15 M WT and α7 KO mice. Upper panels: 4 ×, scale bar 100 μm; Middle panels: stratum oriens (so) and stratum pyramidalis (pyr) of CA1 area, 40 ×; Lower panels: stratum radiatum (sr), 40 ×, scale bar 50 μm. On the right, bar graph showing an increase of neuronal loss in hippocampi from α7 KO (t_(34)_ = 3.938; P < 0.0001). On the bottom, analyses of CA1 layers evidenced a significant loss of neurons in the stratum pyramidalis of α7 KO hippocampi (t_(34)_ = 3.617; P = 0.001). No differences were detected in stratum radiatum and stratum oriens. N = 6 sections from 3 animals/genotype. **B)** Representative images of GFAP staining in the hippocampus of 12-15 M WT and α7 KO mice. Upper panels: 4 ×, scale bar 100 μm; Middle panels: so and pyr of CA1 area, 40 ×; Lower panels: sr, 40 ×, scale bar 50 μm. On the right, bar graph showing an increase of GFAP positive cells in hippocampi from α7 KO animals compared to WT (t_(34)_ = 5.874; P < 0.0001). On the bottom, analyses of CA1 layers showed astrocytosis in the three layers: stratum oriens (t_(34)_ = 4.786; P < 0.0001), stratum pyramidalis (t_(34)_ = 4.621; P < 0.0001), and stratum radiatum (t_(34)_ = 4.321; P < 0.0001). N = 6 sections from 3 animals/genotype. ***P < 0.001, ****P < 0.0001. Data expressed as mean ± SEM.

Finally, because either the increase of Aβ and tau as well as the loss of neuronal function influence astrocytes (González-Reyes et al., 2017; Phatnani and Maniatis, 2015), we evaluated GFAP (Glial Fibrillary Acidic Protein) positive astrocytes. Quantification of GFAP-positive cells revealed an increase that was significant in all the hippocampal strata, i.e. *oriens*, *pyramidalis* and *radiatum*, in hippocampi from α7 KO compared to WT mice (Figure 5B).

Taken together, these findings suggest that α7 KO present a decrease of neurons and astrocytosis in the hippocampus.

## 4. Discussion

In this work we showed that genetic deletion of α7nAChRs is sufficient to induce an AD-like pathology characterized by synaptic plasticity and memory impairment, Aβ and tau neuropathology, neuronal loss, and astrocytosis.

We used the α7 KO mouse model that does not overproduce Aβ due to manipulations of genes directly involved in its production (i.e., APP or presenilins). This allowed us to avoid one of the main limitations of research in the AD field, which is the use of models inspired by rare forms of inherited early onset Familial Alzheimer’s disease (FAD) characterized by a genetic-driven rise of Aβ production (Hardy et al., 1998). Indeed, FAD only accounts for the 2-3% of AD cases (Qiu et al., 2009), whereas the prevalent form of dementia is sporadic AD, which affects the elderly and it is not associated with genetic mutations directly leading to an increase of Aβ burden (Herrup, 2015).

We first focused on synaptic plasticity, whose disruption is thought to be the early pathogenetic event in AD (Selkoe, 2002). α7 KO mice presented an age-dependent impairment of short- and long-term plasticity, as indicated by the increase of PPF and the reduction of LTP, resulting in memory loss. This is consistent with previous studies indicating that, at physiological concentrations, Aβ requires α7nAChRs to sustain synaptic functions (Lawrence et al., 2014; Mura et al., 2012; Puzzo et al., 2011, 2008), both at pre-synaptic level, where it enhances release probability (Gulisano et al., 2019; Koppensteiner et al., 2016; Lazarevic et al., 2017), and at post-synaptic level, where it is needed for long-lasting LTP and memory formation (Garcia-Osta and Alberini, 2009; Gulisano et al., 2019; Morley et al., 2010; Palmeri et al., 2017; Puzzo et al., 2011).

Considering that α7nAChRs mediate the raise of pre-synaptic intracellular Ca^2+^ levels during neuronal activity, thus modulating glutamate release, synaptic transmission, and cognitive function (Picciotto, 2000), it was crucial to exclude that the observed α7 KO phenotype was exclusively due to the failure of cholinergic transmission. Our finding that the impairment of synaptic plasticity and memory is paralleled by the increase of APP expression and Aβ levels suggests that the loss of α7nAChRs might trigger a chain of events through a negative feedback mechanism aimed at restoring calcium entry inside neurons by stimulating Aβ production. Even if we cannot exclude other mechanisms, our hypothesis is supported by extensive literature demonstrating a tight link between α7nAChRs and Aβ (Oz et al., 2013). Aβ and α7nAChRs co-localized inside neurons or at extracellular level in senile plaques (Nagele et al., 2002; Wevers et al., 1999) and high Aβ levels induced α7nAChR alterations (Dineley et al., 2002, 2001; Dougherty et al., 2003; Li et al., 2011; Liu et al., 2001), whereas α7nAChR activation reduced Aβ synthesis and protected against its toxic effects (Buckingham et al., 2009). Furthermore, the lack of α7nAChRs accelerated the pathology in the Tg2576 mouse model of AD ensuing decrease of hippocampal ChAT activity paralleled by a pronounced loss of pyramidal neurons (Hernandez et al., 2010). In 3×Tg mice a reduction of α7nAChRs was found in the same brain regions where intraneuronal Aβ42 accumulation occurred, determining cognitive deficits (Oddo et al., 2005). Interestingly, an increase of α7nAChR-specific antibodies, able to induce Aβ accumulation and memory impairment in animal models (Lykhmus et al., 2015), has been found in plasma samples of early-onset AD patients (Koval et al., 2011).

The increase of APP we found in α7 KO mice raises another series of considerations. Its occurrence in adult age might explain why α7 KO mice do not present a peculiar phenotype until the age of 12 months. Even if it is beyond the scope of this work, we can speculate that in an initial phase other nAChR subtypes, i.e., α4β2nAChRs, might compensate for the absence of α7nAChRs, a common phenomenon in genetically modified animal models. Moreover, it is conceivable that homeostatic changes occur overtime until a critical stage, when compensation is no longer possible. This trend mirrors the course of the disease since synaptic disruption is thought to begin long before the clinical manifestation. Nevertheless, even if in α7 KO mice the increase of APP is aimed at restoring Aβ function, it eventually leads to a vicious cycle in which Aβ levels reach high concentrations becoming extremely neurotoxic.

In this context, APP might exert a double role as it acts as Aβ precursor and cell surface receptor (Deyts et al., 2016) able to bind Aβ and tau (Fogel et al., 2014; Lorenzo et al., 2000; Shaked et al., 2006; Takahashi et al., 2015; Van Nostrand et al., 2002). APP enhances tau phosphorylation (Greenberg et al., 1994) and serves as a common target for extracellular oligomers of Aβ and tau to enter neurons and impair LTP and memory (Puzzo et al., 2017; Wang et al., 2017). Hence, increased APP expression might contribute to worsening the course of the disease with different mechanisms.

As for the interplay between α7nAChRs and tau in AD, results are conflicting (Rubio et al., 2006). Although some works have shown that the increase of α7nAChRs stimulates tau phosphorylation (Ren et al., 2007; Wang et al., 2003), most studies showed that a reduction of α7nAChRs is concomitant with tau hyperphosphorylation in brains of AD patients or animal models (Wu et al., 2010). Here, we found that α7nAChRs deletion induced an increased expression of tau phosphorylated at Thr 205, Ser 199 and Ser 396, residues known to be involved in AD onset and progression. In particular, pTau at Thr 205 seemed to be involved in tau spreading, as demonstrated in Tg/hTau mice injected with tau (Miao et al., 2019) and, together with pTau at Ser 199, has been correlated with Braak stage V/VI in patients (Neddens et al., 2018). As for pTau at Ser 396, it is considered a key marker of tau hyperphosphorylation since it increases in CA1 pyramidal neurons of AD patients and is crucial for PHFs formation (Furcila et al., 2018; Mondragón-Rodríguez et al., 2014). Consistently, here we found an increase of its expression by WB and PHF-1 immunoreactivity at hippocampal level.

These modifications of tau phosphorylation were accompanied by a concomitant decrease of GSK-3β phosphorylated at Ser 9, a residue involved in the auto-inhibitory regulation of the kinase. In line with our results, α7nAchR agonists reduced tau phosphorylation *in vitro* and *in vivo* by increasing GSK-3β activity in mouse models of AD and hypothermia-induced tau hyperphosphorylation (Bitner et al., 2009; Hu et al., 2008), effects that are reversed by selective α7nAchR antagonists (Hu et al., 2008).

Another finding of our study is that α7 KO mice exhibited an increase of APP and Aβ levels but no senile plaques. This is in line with several studies demonstrating that soluble oligomers increase in the early stages of AD and appear more toxic than insoluble aggregates [for a review see (Selkoe and Hardy, 2016)]. Consistently, in animal models, low molecular weight Aβ oligomers (dimers), even though unable to initiate plaque formation (Müller-Schiffmann et al., 2016), induce synaptic dysfunction and trigger the AD cascade (Kawarabayashi, 2004; McDonald et al., 2015, 2010; Shankar et al., 2008). Studies performed on the arcAβ mice, carrying the Swedish and the Arctic mutations, confirmed that insoluble Aβ deposits are not needed to initiate the cascade of events leading to AD as the impairment of synaptic plasticity and memory occurs before plaques formation (Knobloch et al., 2007). However, we cannot exclude that in our study the absence of plaques might be due to a lower propensity of murine Aβ to form insoluble deposits, as evidenced in previous studies (Daniela Puzzo et al., 2015; Puzzo et al., 2014b).

The controversial role of Aβ deposition and its poor correlation with AD symptoms is also supported by several observations in humans showing that AD patients can manifest dementia without Aβ deposits and, conversely, plaques might be present in cognitively intact elderly subjects (Arriagada et al., 1992; Chételat et al., 2013; Delaère et al., 1990; Driscoll et al., 2006; Iacono et al., 2009; Katzman et al., 1988; Sloane et al., 1997; Zolochevska and Taglialatela, 2016).

On the contrary, tau hyperphosphorylation leading to PHFs and NFTs formation is highly related to cognitive impairment in AD (Nelson et al., 2012), as it also contributes to functional and structural alterations of pyramidal neurons (Merino-Serrais et al., 2011). In this manuscript we have shown that α7 KO mice at 12 months presented all these tau-related pathologic signs.

Dementia has also been strongly correlated with the degree of neuronal loss especially in the hippocampus and neocortex in humans (Donev et al., 2009). However, AD mouse models do not always mimic this aspect of the disease (Wirths and Bayer, 2010). Here, NeuN experiments have demonstrated a reduction of neuronal number in hippocampi from α7 KO mice.

Finally, we have documented the presence of astrocytosis in α7 KO hippocampi, in line with previous reports indicating its amplification in AD (Ceyzériat et al., 2018; González-Reyes et al., 2017). Astrocytes are involved in Aβ metabolism and function as they overexpress β-secretases and stimulate Aβ production (Rossner et al., 2005) but also participate in Aβ clearance in physiological conditions (Mulder et al., 2012; Ries and Sastre, 2016). Furthermore, they are involved in the activation of intracellular signaling leading to tau hyperphosphorylation (Chiarini et al., 2017). Several reports have also highlighted a crosstalk between astrocytes and the cholinergic system (Pirttimaki et al., 2013). Indeed, septo-hippocampal lesions of cholinergic fibers induce astrocytosis and β-secretase overexpression (Hartlage-Rübsamen et al., 2003), whereas activation of α7nAChRs by physiological and pathological concentrations of Aβ triggers Ca^2+^ elevations and glutamate release from astrocytes (Lee et al., 2014; Pirttimaki et al., 2013).

In conclusion, here we have demonstrated that α7nAChRs malfunction might be upstream the increase of Aβ and tau in the cascade of events leading to AD, supporting the hypothesis that if Aβ lacks its endogenous receptor, a negative feedback mechanism is triggered to overcome the failure of its physiological function. Even if AD is a multifactorial disease and α7nAChRs malfunction might not be the only etiopathological factor, our findings contribute to understand why ACh-tailored therapies have a time-limited efficacy. In fact, the increase of ACh in the synaptic cleft by cholinesterase inhibitors or the use of AChR agonists might counteract the disease only for a brief period of time after which they might even be responsible for an exhaustion of the cholinergic system. Most importantly, considering the plethora of evidence supporting the importance of Aβ in synaptic function, anti-Aβ therapies might represent a paradox. In fact, they aim at decreasing the level of a physiological protein whose increase might have a compensatory significance. To summarize with a provocative but enlightening conceptual comparison, it would be as administering anti-insulin drugs in type II diabetes, where hyperinsulinemia is the mere consequence of a compensatory mechanism aimed at counterbalancing the lack of its function caused by receptors resistance.

Even if further studies are needed to better delineate Aβ, tau and α7nAChRs crosstalk, our data suggest that the role of Aβ in AD needs to be reassessed, taking into account mechanisms underlying the transition from physiology to pathology to ensure a novel, safe and rational approach to patients. To this end, α7 KO mice might represent an interesting model to evaluate the cascade of events leading to the increase of Aβ without exploiting FAD human genes.

## Acknowledgements

We thank Peter Davies (Albert Einstein College of Medicine, Bronx, NY, USA) for providing the PHF-1 antibody.

## Author contributions

M.R.T. contributed to conceptualization, behavioral and imaging studies, writing of the original draft; D.D.L.P. performed western blot experiments and contributed to the writing of the manuscript; M.M. contributed to conceptualization, execution and supervision of imaging studies; W.G. contributed to conceptualization, electrophysiological experiments, writing of the original draft; O.A. contributed to the conceptualization, writing, review and editing of the manuscript; CG and FC contributed to the conceptualization, formal analysis of the data, writing, review and editing of the manuscript and provided resources; DP contributed to the conceptualization, formal analysis, visualization, writing of the original draft, supervised the whole work, administered the project. All authors discussed results and commented on the manuscript.

